# Tissue-specific and spatially dependent metabolic signatures perturbed by injury in skeletally mature male and female mice

**DOI:** 10.1101/2024.09.30.615873

**Authors:** Hope D. Welhaven, Avery H. Welfley, Priyanka P. Brahmachary, Donald F. Smith, Brian Bothner, Ronald K. June

## Abstract

Joint injury is a risk factor for post-traumatic osteoarthritis. However, metabolic and microarchitectural changes within the joint post-injury in both sexes remain unexplored. This study identified tissue-specific and spatially-dependent metabolic signatures in male and female mice using matrix-assisted laser desorption ionization-mass spectrometry imaging (MALDI-MSI) and LC-MS metabolomics. Male and female C57Bl/6J mice were subjected to non-invasive joint injury. Eight days post-injury, serum, synovial fluid, and whole joints were collected for metabolomics. Analyses compared between injured, contralateral, and naïve mice, revealing local and systemic responses. Data indicate sex influences metabolic profiles across all tissues, particularly amino acid, purine, and pyrimidine metabolism. MALDI-MSI generated 2D ion images of bone, the joint interface, and bone marrow, highlighting increased lipid species in injured limbs, suggesting physiological changes across injured joints at metabolic and spatial levels. Together, these findings reveal significant metabolic changes after injury, with notable sex differences.

**Significance statement:** Osteoarthritis, the leading cause of disability worldwide, disproportionately affects females with sex being one of the strongest predictors of disease. This disparity is partly driven by sex-specific differences in injury susceptibility, increasing the likelihood of traumatic injury to the anterior cruciate ligament (ACL), other ligaments, and menisci. Using a non-invasive injury model, we demonstrate that injury perturbs the local joint environment and has systemic effects in a sex-specific manner. Furthermore, by leveraging matrix-assisted laser desorption ionization-mass spectrometry imaging of the joint, we provide new insight into the composition of osteochondral tissue at the metabolite level. These sexually dimorphic metabolic responses to joint injury advance current understanding of the complex sexual dimorphism in OA pathogenesis providing a foundation for targeted therapeutic strategies and improved patient outcomes for female patients.

## Introduction

Post-traumatic osteoarthritis (PTOA) accounts for 12% of osteoarthritis (OA) cases, resulting in at least 5.6 million cases annually[1, 2]. The most prevalent risk factor for PTOA is joint injury. 250,000 anterior cruciate ligament (ACL) injuries occur annually, commonly in young individuals aged 16-24. Upwards of 50% of patients will develop PTOA within 10-20 years of injury[3–6]. Furthermore, female sex is a risk factor for joint injury: young female athletes are 2-8 times more likely to sustain a traumatic knee injury requiring surgical repair compared to males[7, 8]. Later in life, females are more likely to develop OA and typically experience greater symptom severity[9, 10]. Sexual dimorphism in PTOA development may partially result from hormonal and anatomical differences including females have wider pelves, smaller femurs, different muscle angles, and physically smaller ACLs[11, 12]. Despite these empirical sex differences, many studies fail to include both female and male subjects.

While injury and sex are well-known PTOA risk factors, underlying mechanisms and changes in joint metabolism and microarchitecture following injury remain incompletely understood. Metabolomics–the study of small molecule intermediates called metabolites– captures the physiological and metabolic status of the joint. While synovial fluid (SF) and serum are assessed post-injury in human and mouse models[13–17], metabolomics can also be applied to whole joint tissue to characterize metabolic changes occurring in bone and cartilage post-injury. This approach combined with spatial imaging through matrix-assisted laser desorption ionization-mass spectrometry imaging (MALDI-MSI) provides critical unknown spatial context of osteochondral tissues at the metabolite level. Few studies report spatial data involving musculoskeletal tissues such as cartilage[21, 22] and synovium[23]. However, no study to date has utilized MALDI-MSI to characterize molecular changes between male and female injured and naïve mice.

Therefore, the first objective of this study was to characterize metabolomic differences within and across whole joints, SF, and serum from injured and naïve male and female mice using liquid chromatography-mass spectrometry (LC-MS) metabolomics. Investigating differences in metabolism in different tissues and between injured, contralateral, and naïve limbs is important to shed light on both local and systemic metabolic responses following injury. The second objective was to spatially locate and identify osteochondral metabolites using MALDI-MSI. By combining untargeted metabolomic profiling and MALDI-MSI, we can pinpoint the origin of metabolic and pathological shifts and gain a better understanding of the effects of injury and sex on the joint as a whole.

## Results

### Injury Perturbs the Metabolome Systemically Across Injured and Naive Whole Joints, Synovial Fluid, and Serum

In total, 2,769 metabolite features were detected across all samples (n=92). Whole joints, SF, and serum were assessed among all samples and injured samples only revealing distinct metabolomic profiles between tissues and fluids as showcased by PLS-DA (Fig. S1A-B). ANOVA found 2,264 and 1,891 metabolite features that were significantly dysregulated across all samples and only injured samples, respectively (Fig. S1C-D). Next, we assessed metabolic patterns associated with injury within each sample type. PLS-DA assessed metabolomic profiles by injury status across whole joint, SF, and serum (Fig. 1A-C). Fold change analysis identified populations of metabolite features driving these differences between injury groups, which were subjected to pathway analysis (Fig. 1D-F). Comparing injured and naïve whole joints, 7 pathways were dysregulated. Pantothenate/CoA biosynthesis and histidine metabolism were highest in injured whole joints (Table S1A). In SF, 12 pathways were dysregulated: arginine and proline metabolism, lysine degradation, and glutathione metabolism were highest in injured SF (Table S1B). Features highest in serum from injured mice mapped to various amino acid pathways, including arginine and proline metabolism, nucleotide pathways, vitamin, and glutathione metabolism (Table S1C). Next, metabolic indicators of injury were assessed using volcano plot analyses, comparing features differentially regulated across injured and naïve samples (Fig. S2A-C). Cross-referencing these features with LC-MS/MS data identified 3,4-Dimethyl-5-pentyl-2-furanundecanoic acid as differentially regulated between naïve and injured whole joints, while numerous amino acid metabolites, such as D/L-glutamine, were higher in injured SF compared to naïve (Table S2).

**Figure 1.**
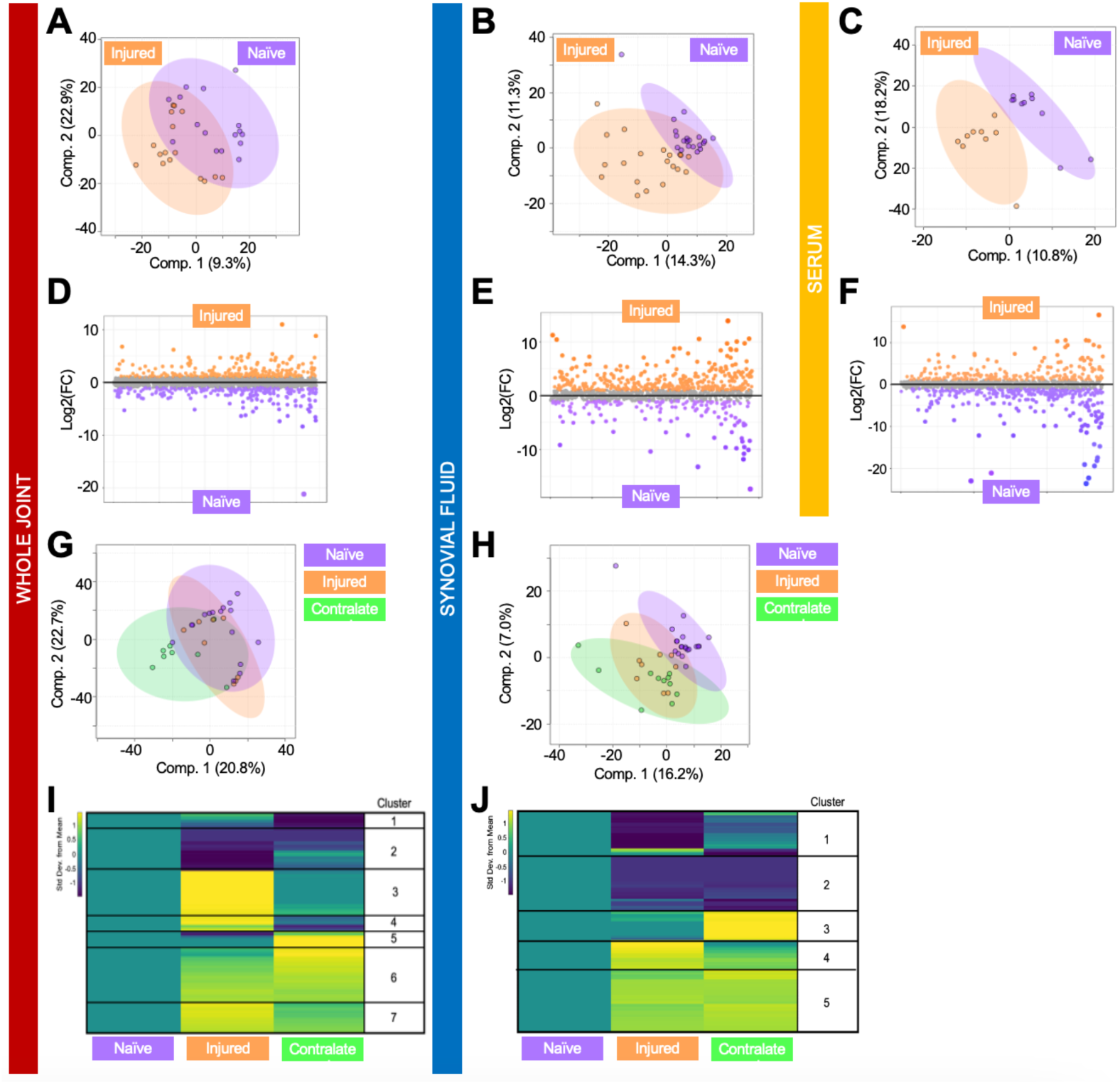
Global metabolomic profiles of whole joints, synovial fluid, and serum are driven by injury status. (A-C) Partial Least Squares-Discriminant Analysis (PLS-DA) finds some overlap between injured and naïve whole joint and synovial fluid and near-perfect separation of injured and naïve serum. (D-F) Fold change analysis distinguished populations of metabolite features driving separation of metabolomic profiles. (D) Specifically, 250 and 291 metabolite features were highest in injured and naïve whole joints, respectively. (E) 373 and 155 metabolite features were highest in injured and naïve synovial fluid, respectively. (F) 386 and 195 features were highest in injured and naïve serum, respectively. Similarly, PLS-DA reveals overlap between injured, contralateral, and naïve (G) whole joints and (H) synovial fluid with injured samples clustering together between contralateral and naïve samples. To pinpoint pathways driving metabolomic differences between limbs with different injury statuses, median intensity heatmap analyses where injured and contralateral limbs were normalized to naïve limbs were performed. Clusters of co-regulated metabolite features within (I) whole joint and (J) synovial fluid samples were subjected to pathway analyses to identify biological pathways that differ in regulation across limbs in both whole joint and synovial fluid samples. Combined, data provide strong evidence of distinct metabolomic regulation associated with injury status. Columns represent limbs (naïve, injured, contralateral) and rows represent metabolite features. Cooler and warmer colors indicate lower and higher metabolite abundance relative to the mean, respectively. The colors in A-J correspond to: purple = naïve, orange = injured, green = contralateral whole joint; sample types-red = whole joint, blue = synovial fluid, yellow = serum.

Next, metabolic differences associated with injured, contralateral, and naïve whole joints and SF were examined. PLS-DA analysis of all three joint types revealed overlap, where the metabolome of injured whole joints and SF resides between contralateral and naïve whole joints and SF (Fig. 1G, H), suggesting metabolic differences associated with injury at the whole joint and SF levels are observed in the contralateral limb. Metabolite features that are co-regulated and differentially expressed across whole joints and SF were clustered using heatmaps of median metabolite feature intensities, which were then subjected to pathway analyses (Fig. 1I, J). Comparing injured, contralateral, and naïve whole joints 22 pathways were detected and 24 among SF (Table S3, S4). A handful of pathways were detected in both heatmaps. Amino acid pathways – alanine, aspartate, and glutamate metabolism; arginine and proline metabolism, lysine degradation – were higher in injured and contralateral whole joints and lower in these same groups in SF, compared to naive controls. Notably, glutathione metabolism was consistently highest in injured whole joints and SF compared to contralateral and naïve controls. Purine and pyrimidine metabolic pathways also displayed a similar pattern where these pathways were highest in injured whole joints and SF. These systemic injury-induced effects across limbs in whole joints and SF were examined through pairwise comparisons and are discussed in detail in the supplemental results (Figs. S3-4, Tables S5-6). Notably, unloaded contralateral limbs resembled injured more than naïve limbs, further supporting that injury perturbs beyond the site of injury, and instead has clear systemic effects.

### Sex Influences Metabolomic Profiles Across Tissues of Injured and Naive Mice

To examine the effects of sex and its interactions with injury, pairwise comparisons were performed. First, general sex differences were examined in whole joints (Fig. S5), revealing distinct sex-dependent metabolomic profiles. Then, whole joints from injured males and females were assessed using PLS-DA finding clear separation of mice within their respective cohorts. Differences due to injury are evident when comparing whole joints from injured and naïve females and from injured and naïve males as seen by minimal PLS-DA overlap (Fig. 2A-C). This same pattern was evident in both SF (Fig. 2G-I) and serum (Fig. 2M-O), with serum differences being the most drastic among samples. These findings at the molecular level demonstrate that sex-specific differences emerge post-injury, are more pronounced within the serum, and contribute to the mounting evidence indicating the role of sex in PTOA development.

**Figure 2.**
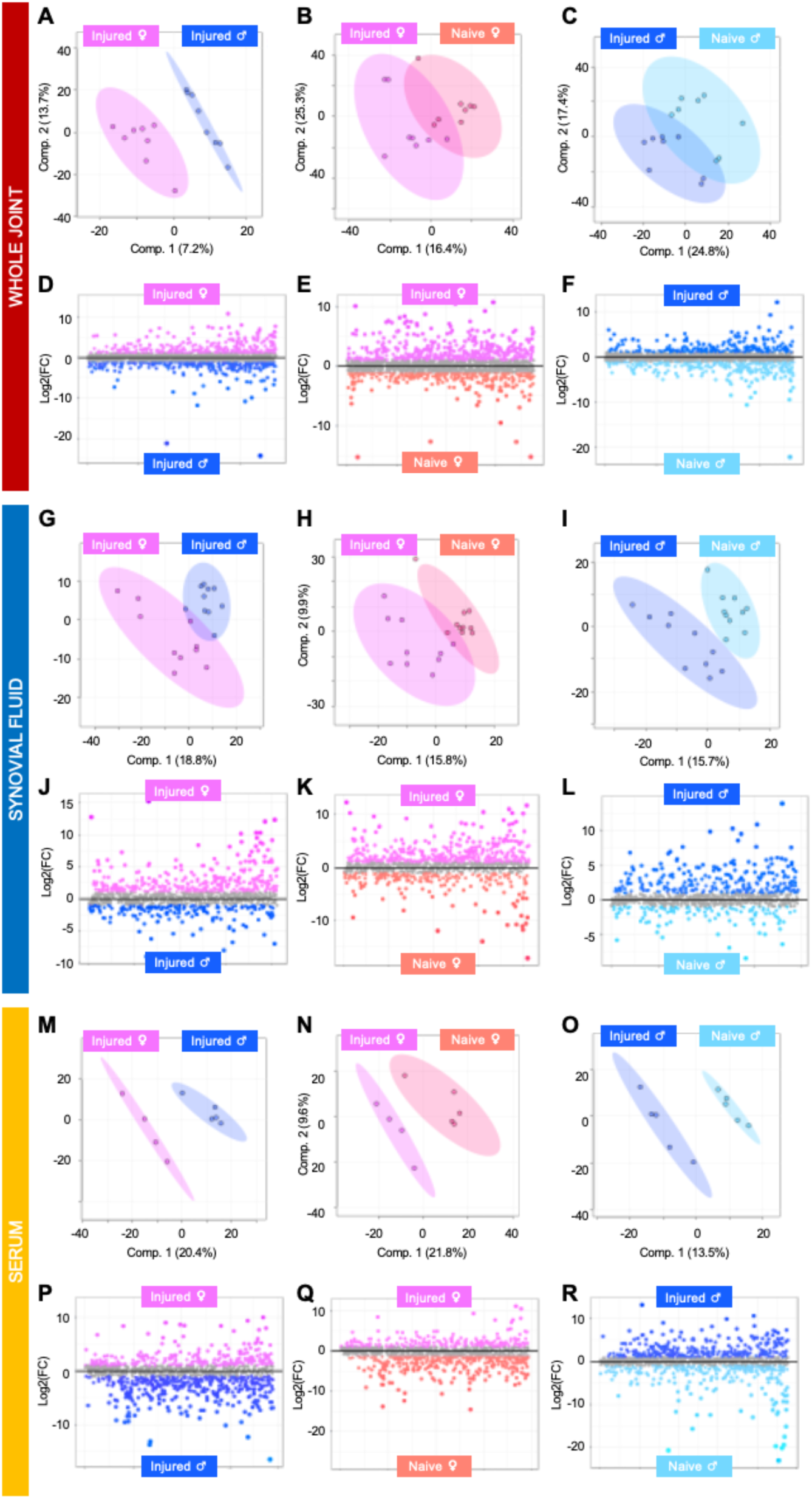
Metabolomic profiles of whole joint, synovial fluid, and serum show sexual dimorphism across injured and naive mice. (A-C) Partial Least Squares-Discriminant Analysis (PLS-DA) finds (A) complete separation of injured whole joints from males and females and minimal overlap when comparing (B) female and (C) male injured and naïve mice. (D-F) Fold change analysis distinguished populations of whole-joint derived metabolite features driving separation of metabolomic profiles. (G-I) PLS-DA finds minimal overlap when comparing (G) injured SF from males and females, (H) female and (I) male injured and naïve mice. (J-L) Fold change analysis identified populations of synovial fluid metabolite features contributing to the separation of mice that differ by sex and injury. (M-O) PLS-DA finds clear separation with no overlap when comparing (M) injured SF from males and females, (N) female and (O) male injured and naïve mice. (P-R) Fold change analysis identified populations of metabolite features driving separation of serum metabolomic profiles. The colors in A-R correspond to: pink = injured females, peach = naïve females, royal blue = injured males, light blue = naïve males. sample types - red = whole joint, blue = synovial fluid, yellow = serum.

Fold change analysis found populations of metabolite features from whole joints, SF, and serum that show distinct injury- and sex-specific pathways with many conserved across all three sample types (Tables S7-9). Alanine, aspartate, and glutamate metabolism were detected in whole joints of injured males and SF and serum of both injured males and females. Arginine biosynthesis was detected among whole joints from males; however, it was also detected in serum from injured females and SF from injured males and females. Beta-alanine, glyoxylate, and dicarboxylate metabolism were detected across all male naïve tissues. Cysteine and methionine metabolism was detected in both whole joint and SF from injured females. Interestingly, lysine degradation and pyrimidine metabolism were detected across all three sample types in injured females. Notably, glycerophospholipid metabolism was continually associated with injury, and was highest in samples from injured males at the whole joint and serum levels, whereas it was highest in SF from injured females.

To investigate metabolic indicators of both injury and sex, volcano plot analysis was performed comparing whole joints, SF, and serum from injured and naïve males and females. This identified sex- and injury-specific metabolites, including those linked to terpenoid backbone biosynthesis – (6R)-6-(L-Erythro-1,2-Dihydroxypropyl)-5,6,7,8-tetrahydro-4a-hydroxypterin in females and Sterebin E in males – in whole joints. Additionally, 1/3-Methylhistamine and valine showed similar patterns across SF and serum, with valine associated with males, and highest in injured males. Notably, 1/3-methyhistidine was associated with injury, regardless of sex, and was highest in SF from injured females (Figure S6, Table S2).

### MALDI-MSI Spatially Locates and Detects Differences in Osteochondral Metabolites

MALDI-MSI was used to characterize metabolites that are tissue-specific in the whole joint. A novel protocol for sample preparation, matrix application, and instrumentation was developed to examine spatial changes. Using unsupervised segmentation, discriminating features from different spatial areas of the joint were clustered together. When comparing spatial distributions of ions belonging to the bone and joint interface against the bone marrow, numerous features had area under the curve (AUC) values greater than 0.6 (n=37) and 0.8 (n=21), demonstrating tissue-dependent metabolite signatures. Using LC-MS/MS-derived metabolite identifications from whole joint samples, a handful of these metabolites with unique spatial patterns were putatively identified. Those with notable spatial patterns in bone included alpha-carboxy-delta-decalactone (215.20 m/z, AUC = 0.838), hydroxyprolyl-isoleucine (245.11 m/z, AUC = 0.859), and C_36_H_38_O_7_ (629.61 m/z, AUC = 0.829) (Fig. 3, Fig. S7A, Table S10A).

**Figure 3.**
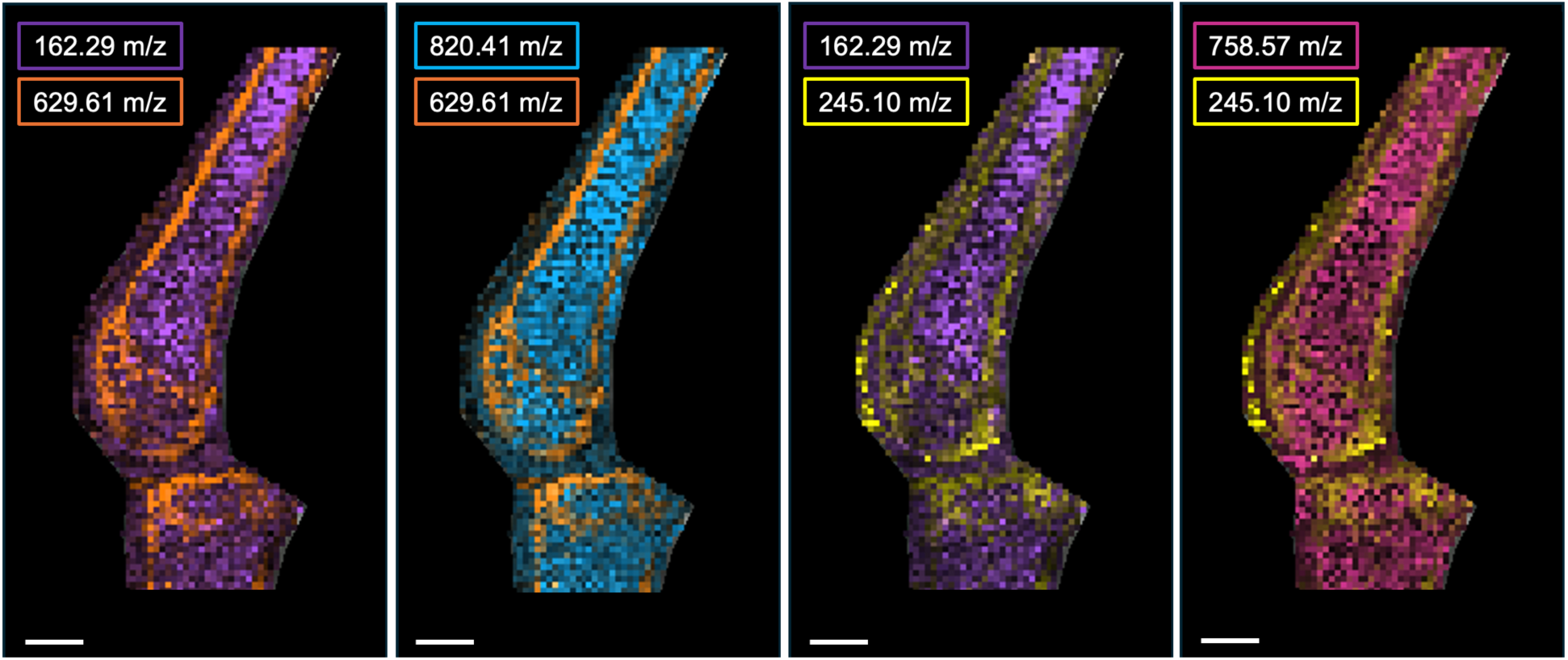
MALDI-MSI combined ion images from sagittal whole joint sections. Heatmap analysis of putatively identified molecular species across various tissue structures. Spatial resolution = 100 um. Scale bar = 1 mm. Interval width = 0.35 Da. Colors in panels from left to right: L-carnitine (162.29 m/z, purple), C_36_H_38_O_7_ (629.61 m/z, orange), (820.41m/z, blue), hydroxyprolyl-isoleucine (245.10 m/z, yellow), and lipid species 18:2/16:0 (758.57 m/z, pink).

Conversely, metabolites with notable patterns among bone marrow included carnitine (162.29 m/z, AUC = 0.821) and various phosphatidylcholine lipid species (34 carbons, 2 double bonds – 758.57 m/z, AUC = 0.731; 34 carbons bonds, 3 double bonds – 778.59 m/z, AUC = 0.856; 39 carbons, 6 double bonds – 820.41 m/z, AUC = 0.866) (Fig. 3, Fig. S7B, Table S10A). Many metabolites displayed unique spatial patterns between bone, the joint interface, and bone marrow but were unable to be identified. However, many of these were within the common lipid m/z range (400-800 m/z) (Fig. S8, Table S10B).

To investigate injury-associated spatial patterns, ion images from medial and lateral aspects of both injured and contralateral mouse joints were examined. In an injured joint, the metabolite feature 544.41 m/z and the putatively identified metabolite–a PC with 36 carbons, 6 double bonds 878.54 m/z–display notable injury-associated patterns between medial and lateral sagittal sections where they are more abundant in the injured joint compared to the contralateral joint (Fig. S9). Sex-associated patterns were not examined at length due to the limited number of joints allocated for MALDI-MSI (n = 1 mouse/group, n = 2 males, 2 females, n = 4 mice total).

## Discussion

To our knowledge, this is the first study to comprehensively examine structural and metabolic responses following injury both locally (whole joints and SF) and systemically (serum) across males and females using LC-MS metabolomic profiling and MALDI-MSI. Our findings reveal significant metabolic and pathologic shifts following joint injury, with discernable sex-specific associations. By uncovering novel metabolic differences linked to these key PTOA risk factors across multiple sample types and biological scales, our study advances the understanding of post-injury responses beyond the joint. These insights provide critical, and previously unknown, context on the joint-specific metabolic and systemic changes that shape disease progression.

### Effects of injury across whole joints, synovial fluid, and serum

Distinct metabolomic profiles of whole joints, SF, and serum reveal acute changes in response to injury. Various amino acid pathways exhibit differential regulation across tissues with many overlapping between tissues. Lysine degradation was differentially regulated across injured, contralateral, and naive whole joints while being downregulated in SF contralateral and injured limbs (versus naïve). This may relate to collagen, where hydroxylation of lysine residues is crucial for structural integrity crosslinking [25]. In bone, collagen is a major component of the structural organic matrix. Alteration of lysyl hydroxylase in osteoblasts yields defective cross-linking, fibrillogenesis, and matrix mineralization[25–27] underscoring the importance of lysine hydroxylation in bone quality. In SF, lysine metabolism decreases as OA progresses[28], suggesting this pathway could represent acute joint changes following injury that could be monitored during disease progression.

Arginine and proline metabolism was upregulated in naïve whole joints and in injured SF. Like lysine, proline can strengthen collagen crosslinks through hydroxyproline modification. Arginine has anti-inflammatory effects and decreases as OA develops[29, 30]. Arginine and proline metabolism increase in rabbit SF post-ACL injury[31], in human SF after knee injuries[32], and in mouse serum 1-day post-ACL injury[15]. Combined, this suggest a short-term protective mechanism, or cascade of pathways to enhance collagen production post-injury.

Phenylalanine, tyrosine, and tryptophan biosynthesis exhibited differential regulation between injured, contralateral, and naïve limbs in whole joint samples and SF. While higher in injured and contralateral whole joints (versus naïve), these pathways showed the opposite trend in SF. Calcium-sensing receptors preferentially bind these aromatic amino acids, leading to a rise in intracellular calcium and modulation of bone turnover[33, 34]. Moreover, phenylalanine and tyrosine metabolism are associated with the sclerosis of subchondral bone in OA[35]. In a noninvasive mouse injury model, tryptophan metabolism was higher in mouse SF 7 days post-injury compared to naïve controls. Tryptophan has been noted as a promising OA biomarker with decreases as disease progresses[36, 37]. Our detection of this pathway in both injured and contralateral whole joints is novel, suggesting a systemic response in both injured and contralateral whole joints. Moreover, these results hold promise as amino acids play a role in the response to acute injury[38], are detected post-injury across mammalian models[15, 16, 31], and change concentration with disease progression.

### Sexual dimorphism of injury among the metabolome of whole joints, synovial fluid, and serum

Considering both injury and sex, purine and pyrimidine metabolism—used for nucleotide synthesis—were dysregulated. Purine metabolism was consistently detected across all tissues from injured males, particularly in serum, which likely results from increased uric acid–an end product of purine metabolism–in male[39] (because female hormones decrease uric acid levels[40]). Conversely, pyrimidine metabolism was dominant in tissues from injured females, necessitating further research to understand its sexual dimorphic regulation.

Dysregulated glycine, serine, and threonine metabolism was detected in serum and SF in injured males and females. This pathway was notably perturbed among female sheep post-ACL injury, with serine suggested as a biomarker for early degenerative changes[14]. Serine, a glucogenic amino acid, influences adenosine monophosphate kinase (AMPK), which acts as a key “energy sensor” that maintains energy homeostasis and promotes ATP synthesis via serine/threonine phosphorylation[41–43]. Enzymes like AMPK are differentially influenced by circulating sex hormones like estrogen, which can bind to estrogen receptor beta, triggering downstream metabolic cascades to generate ATP[44, 45]. The influence of circulating sex hormones, like estrogen, on AMPK activity may affect energy metabolism post-injury, warranting further investigation into sexual dimorphic patterns.

Cysteine and methionine metabolism was detected in whole joints and SF from injured females but also in serum from injured males. Histidine metabolism was most dysregulated in injured males in serum and whole joints and in injured females SF. These three amino acids relate to matrix metalloproteinase (MMP) regulation of tissue remodeling and degradation of extracellular matrix proteins, cell proliferation, and immune responses[46]. MMP activation is modulated by a cysteine switch, and the catalytic domain of MMPs is regulated by these amino acids because zinc binds to histidines with assistance from conserved methionine sequences[46, 47]. MMP activity is influenced by hormones[48], like estrogen and progesterone, particularly in chondrocytes. Postmenopausal OA chondrocytes cultured with 17B-estradiol found that physiological levels of estrogen suppressed the expression of MMP-1 and that hormone replacements might benefit female OA patients in the early stages of disease[49]. Thus, differential regulation of AMPK and MMP-associated amino acids between males and females post-injury suggests reliance on different metabolic pools, mechanisms, and biofuels to meet energy demands and maintain matrix properties following joint injury.

### MALDI-MSI Provides New Insight into Composition of Osteochondral Tissue at the Metabolite Level

To our knowledge, this is the first study to provide critical unknown spatial context of osteochondral tissues at the metabolite level, and after joint injury. We developed an innovative protocol to spatially characterize osteochondral metabolites from whole joints of injured and naïve mice using MALDI-MSI. Few studies have employed MALDI-MSI in musculoskeletal tissues to visualize the spatial distribution of proteins and peptides[20, 23, 28, 50]. Traditionally, histological sectioning and imaging of whole joints—and bone samples in general—require formalin fixation and paraffin embedding to preserve and demineralize the bone; however, this results in removal, cross-linking, and/or delocalization of molecular species, especially lipids[24], leading us to develop this novel protocol to examine spatial metabolite distributions.

Lipid metabolism is increasingly recognized in the development of PTOA and OA. Proteomic and metabolomic studies demonstrate an important relationship between OA and lipid metabolism in samples of SF, cartilage, bone, and circulatory fluids from both humans and mice[21, 28, 51–54]. Employing MALDI-MSI to target small molecules (50-1500 m/z) provides novel insight into joint composition, lipid dynamics, and changes post-injury. We successfully mapped differences in injury-related osteochondral metabolites, where many were lipids. These findings align with metabolomic assessments finding lipid-associated pathways—such as glycerophospholipid metabolism—higher in injured joints compared to naive and contralateral joints. Consistent with our previous analysis of metabolic endotypes of early PTOA in SF[17, 28, 32, 50] and in OA cartilage[54, 55], dysregulated lipid metabolism is linked to both early-stage PTOA and end-stage OA, suggesting similar metabolic shifts occur in the joint at the metabolic and spatial levels. This underscores the need for further research into lipid metabolism’s role in both local and systemic changes following injury and during early PTOA. This two-pronged approach utilizing LC-MS-based metabolomics and MALDI-MSI sheds light on the importance of lipids in joint metabolism, both systemically and intra-articularly. Moving forward, MALDI-MSI across proteins and metabolites can be leveraged and integrated with existing techniques to enhance our understanding of the joint’s post-injury response across multiple tissues.

Our study has limitations. Firstly, we focused on the early response to joint injury where mice were euthanized 8 days post-injury. Additional time points both within the first 7 days post-injury as well as longer[56] may shed light on the trajectory of structural and metabolic changes across serum, SF, and whole joints. Secondly, metabolite extraction protocols and use of HILIC column bias data toward the assessment of polar molecules. Moreover, DHB was used for MALDI-MSI, which is optimal for positive ionization mode. Combined, LC-MS and MALDI-MSI analyses were both conducted in positive ionization mode focusing on polar molecules, thus additional investigation into nonpolar species is warranted.

### Conclusions

The findings of our study, integrating LC-MS-based metabolomics and MALDI-MSI, underscore significant metabolic and pathological shifts following joint injury, with discernible sex-specific associations. The detection of novel differences associated with both injury and sex across serum, SF, and whole joints expands our current understanding of the post-injury response within and beyond the joint. Moreover, these data show that injury drives whole joint pathophysiology across multiple tissues. Further, differences in the SF metabolome compared to naïve animals show that the systemic response extends to both injured and contralateral joints. Comprehensive investigation into spatial and sex-dependent molecular changes driven by injury, at the systemic, joint, and synovial fluid levels, is imperative for a deeper understanding of the effects of injury. Extension of these studies may improve pre-clinical PTOA models and deepen our insights into PTOA development, thereby advancing strategies for prevention and treatment.

## Materials and Methods

### Animals

C57Bl/6J mice (N=20, n = 10 female, n = 10 male) were purchased from Charles River Laboratories at 18 weeks of age and acclimated to the Montana State University (MSU) Animal Research Center for 3 weeks. Mice were housed in cages of 3-5 animals and fed standard chow *ad libitum* (PicoLab Rodent Diet 20, 20% protein). All animal procedures were approved by the MSU IACUC.

### Joint Injury Model and Experimental Design

21-week-old mice were randomly assigned to experimental groups: injured or non-injured. The injury group were subjected to a non-invasive compressive overload model where the ACL is ruptured similar to human ACL tears, an injury associated with degeneration of bone and cartilage (target force = 12N, loading rate = 130 mm/s) [57, 58]. ACL injury was confirmed by laxity tests. After 8 days, mice were euthanized and whole joints, SF, and serum were harvested. Serum samples were obtained by cardiac puncture and prepared as previously described[59]. SF was recovered and then extracted using established protocols[60], with some modifications where Whatman paper (Sigma, WHA1441042) was used to absorb SF. Whole joints were harvested, and all soft tissue was removed.

### Metabolomics

#### Metabolite Extraction and Mass Spectrometry Instrumentation

Whole joints, SF, and serum were extracted and analyzed using validated protocols with slight modifications[32]. Whole joints were disarticulated to separate the tibia and femur where both were trimmed, centrifuged to remove bone marrow, and homogenized in 1 mL of 80:20 methanol:H2O (1200 GenoLyte, Fischer Scientific). This same extraction solvent was added to serum and SF, followed by vortexing. All samples were chilled overnight at -20°C. The next day, samples were removed from -20°C, centrifuged, and supernatant was dried via vacuum concentration. To remove any remaining proteins, lipids, and waxes, dried extracts were resuspended with 250 uL of 1:1 acetonitrile:H2O, vortexed, chilled at -20°C for 30 minutes, and centrifuged again. Supernatant was dried once more, and extracts were prepped for LC-MS using 100 uL of 1:1 acetonitrile:water. Additionally, pooled samples were generated by randomly pooling a total of 50 uL for each tissue type (n=4/tissue). Pooled samples from each tissue type were also pooled for identification purposes. All solvents used were LC-MS grade (Fischer Scientific). Extracted samples and pools were analyzed via LC-MS as previously described[32].

#### Metabolite Profiling and Identification

All LC-MS data—mass-to-charge ratios (m/z), relative metabolite abundance, and retention time—were processed using MSConvert and Progenesis QI (Table S11). Statistical and pathway analyses were performed in MetaboAnalyst (version 6.0)[61]. Significance for statistical and pathway analyses was determined using a FDR-corrected significance level of p < 0.05.

LC-MS/MS data derived from pooled samples were analyzed within Progenesis QI where acquired fragmentation patterns were matched against theoretical fragmentation patterns to identify metabolites[32]. Those identified were matched against populations of LC-MS-based features distinguished by statistical analyses to discover potential metabolic indicators of disease as well as sexually dimorphic metabolites. To minimize false identifications when comparing LC-MS and LC-MS/MS metabolite features and identifications, a tolerance level of 10 parts per million was enforced.

#### Matrix-Assisted Laser Desorption Ionization Imaging Sample Preparation

Whole joints selected for MALDI-MSI (n = 1 mouse/group, n = 2 joints/mouse) were removed from -80°C and prepped according to the novel protocol developed for whole joint sectioning and imaging (Fig. S10). Whole joints were embedded using warm 5% carboxymethylcellulose sodium salt (ThermoFischer, A18105-36) and 10% gelatin (Thermo Scientific, AC611995000)[24, 62]. Note that OCT media is incompatible with mass spectrometry analysis due to the presence of polyethylene glycol, a major ion suppressor.

Medial and lateral aspects of the joint were sectioned sagittally at 8 um thickness in a cryostat set to -30°C (OTF5000 Cryostat, Bright Instrument Co Ltd, Tissue-Tek Accu-Edge 4689 blade). Because whole joints did not undergo formalin fixation or paraffin embedding, Cryofilm 3C 16 UF (SECTION-LAB, Hiroshima, Japan) was used to assist in transferring sections to indium tin oxide (ITO) slides (Delta Technologies, CB-401N, 4-10 Ω/sq, 25 x 50 mm)[63]. Sections were adhered to ITO slides using double-stick tape and then directly stored at -80°C until matrix application.

#### Matrix Application

A sublimation apparatus (Chemglass Life Sciences, CG-3038) was used for matrix application (Fig. S11). Sublimation was chosen over matrix spraying as it forms smaller, more homogenous matrix crystals and lacks liquid—reducing the risk of molecule migration and microcracks in marrow and bone[24]. 300 mg of 2,5-dihydroxybenzoic acid (DHB) (Alfa Aesar, 490-79-9) was uniformly dispersed in the base of the sublimation apparatus, ITO slides were affixed to the flat bottom condenser, and the condenser was filled with tap water (22°C). The two glass compartments of the sublimator were assembled using an O-ring seal and connected to a vacuum pump. Tissue sections on ITO slides were sublimated at 50°C for 3 minutes under 68 mTorr vacuum, resulting in a uniform matrix layer (0.05 mg/cm^2^) that was then recrystallized in a hydration chamber with 300 uL of 5% methanol 0.1% formic acid spotted onto Whatman paper. The chamber was heated in a 37°C oven for 1 minute, then ITO slides were sealed in the chamber and heated at 37°C for 1 minute.

#### MALDI Image Acquisition and Analysis

A Bruker AutoFlex III MALDI Time-of-Flight (TOF) mass spectrometer (Bruker Daltonics) equipped with a MTP Slide Adapater II Imaging Plate (Bruker Daltonics) was used for image acquisition. Due to the size of the sublimator and ITO slides (25 x 50 mm), two small aluminum slide trays were milled to fit ITO slides into the imaging plate. Using a Smartbeam Nd:YAG laser (355 nm) and Bruker FlexImaging, images were collected in positive ionization and TOF modes in the 50-1000 m/z mass range averaging 200 laser shots per pixel with a 100-um lateral resolution (laser power = 30%, range = 90%, offset = 10%). Imaging data collected were analyzed and ion images were generated using Bruker SCiLS lab.

## Author contributions

Conceptualization: HDW, RKJ

Methodology: HDW, AHW, PB, DS, BB, RKJ

Investigation: HDW, AHW, DS

Visualization: HDW, AHW, DS

Supervision: DS, BB, RKJ

Writing—original draft: HDW, AHW

Writing—review & editing: HDW, AHW, PB, DS, BB, RKJ

## Data and materials availability

All data are available in the main text and the supplementary materials. Raw metabolomics data is available in table S11.

## Supplemental Figures

**Figure S1.**
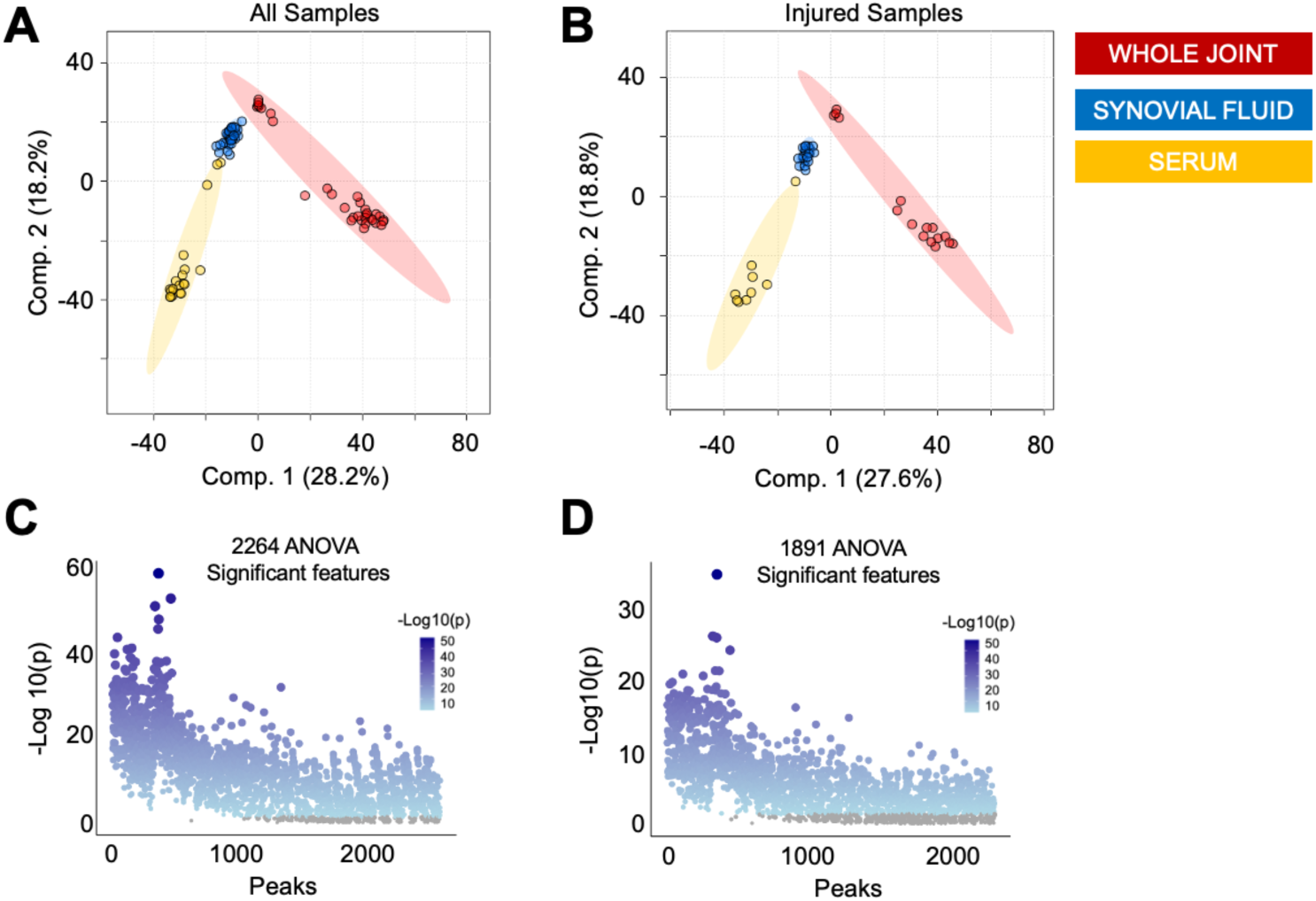
The metabolome is distinct for whole joints, synovial fluid, and serum. Partial Least Squares-Discriminant Analysis (PLS-DA) of (A) all samples and (B) only samples from injured mice displays clear separation of whole joints, synovial fluid, and serum suggesting the metabolome of each tissue is substantially distinct from others. (C) ANOVA analysis identified 2,264 metabolite features that were significantly dysregulated across samples from all three tissue types. (D) When comparing samples from injured mice only, ANOVA analysis identified 1,891 significantly dysregulated features across tissue types. Colors in A-B correspond to: red = whole joint, blue = synovial fluid, yellow = serum.

**Figure S2.**
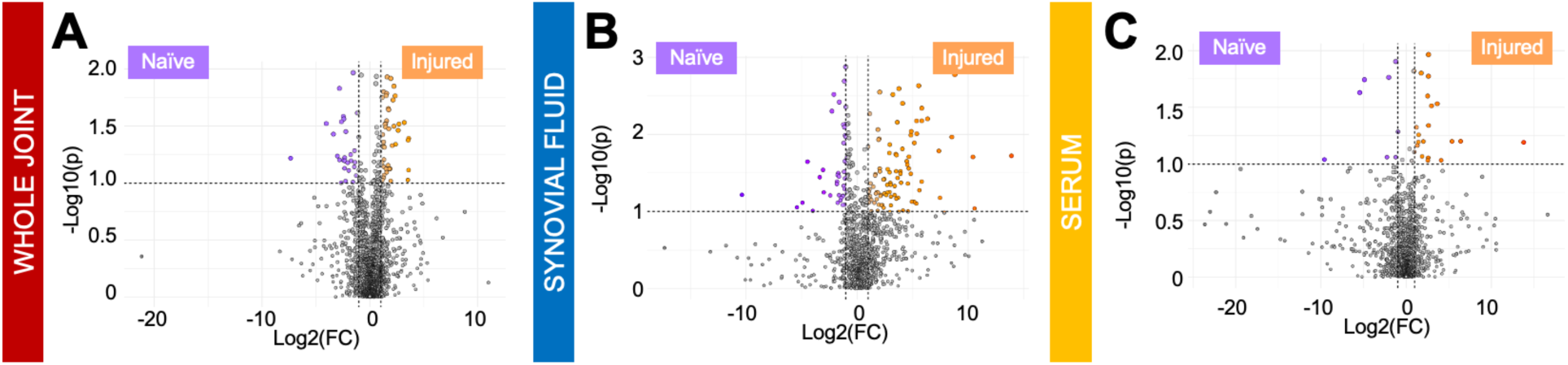
Volcano plot analysis reveals injury-associated metabolites. (A-C) To further examine metabolic differences associated with injury status across tissue types, volcano plot analysis was performed and identified numerous metabolite features that had a fold change > 2, a p-value < 0.05, and were differentially regulated between injured and naïve whole joints (A, n = 70, red), synovial fluid (B, n = 131, blue), and serum (C, n = 26, yellow).

**Figure S3.**
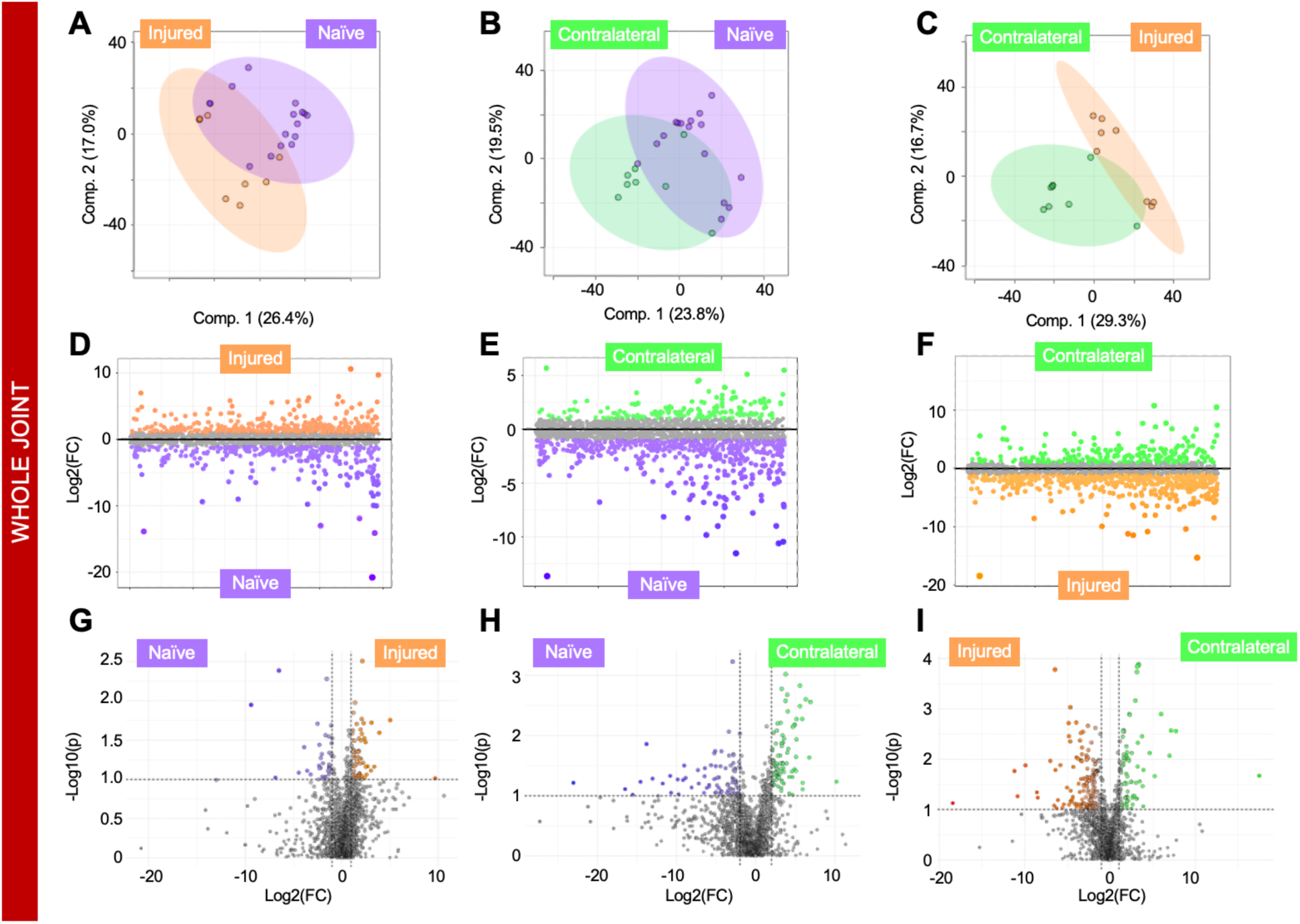
Whole joint metabolome differs by sex and injury. (A-C) Partial Least Squares-Discriminant Analysis (PLS-DA) finds (A) complete separation of injured whole joints from males and females and minimal overlap when comparing (B) female and (C) male injured and naïve mice. (D-F) Fold change analysis distinguished populations of metabolite features driving separation of metabolomic profiles. (D) Specifically, 315 and 314 metabolite features were highest in injured females and males, respectively. (E) 509 and 319 metabolite features were highest in injured females and naïve females, respectively. (F) 242 and 288 features were highest in injured males and naïve males, respectively. (G-I) To further examine metabolic differences associated with injury and sex, volcano plot analysis was performed and identified numerous metabolite features that had a fold change > 2, a p-value < 0.05, and were differentially regulated between injured males and females (G, n = 110), injured and naïve females (H, n = 158), and injured and naïve males (I, n = 32). The colors in A-I correspond to: pink = injured females, peach = naïve females, royal blue = injured males, light blue = naïve males.

**Figure S4.**
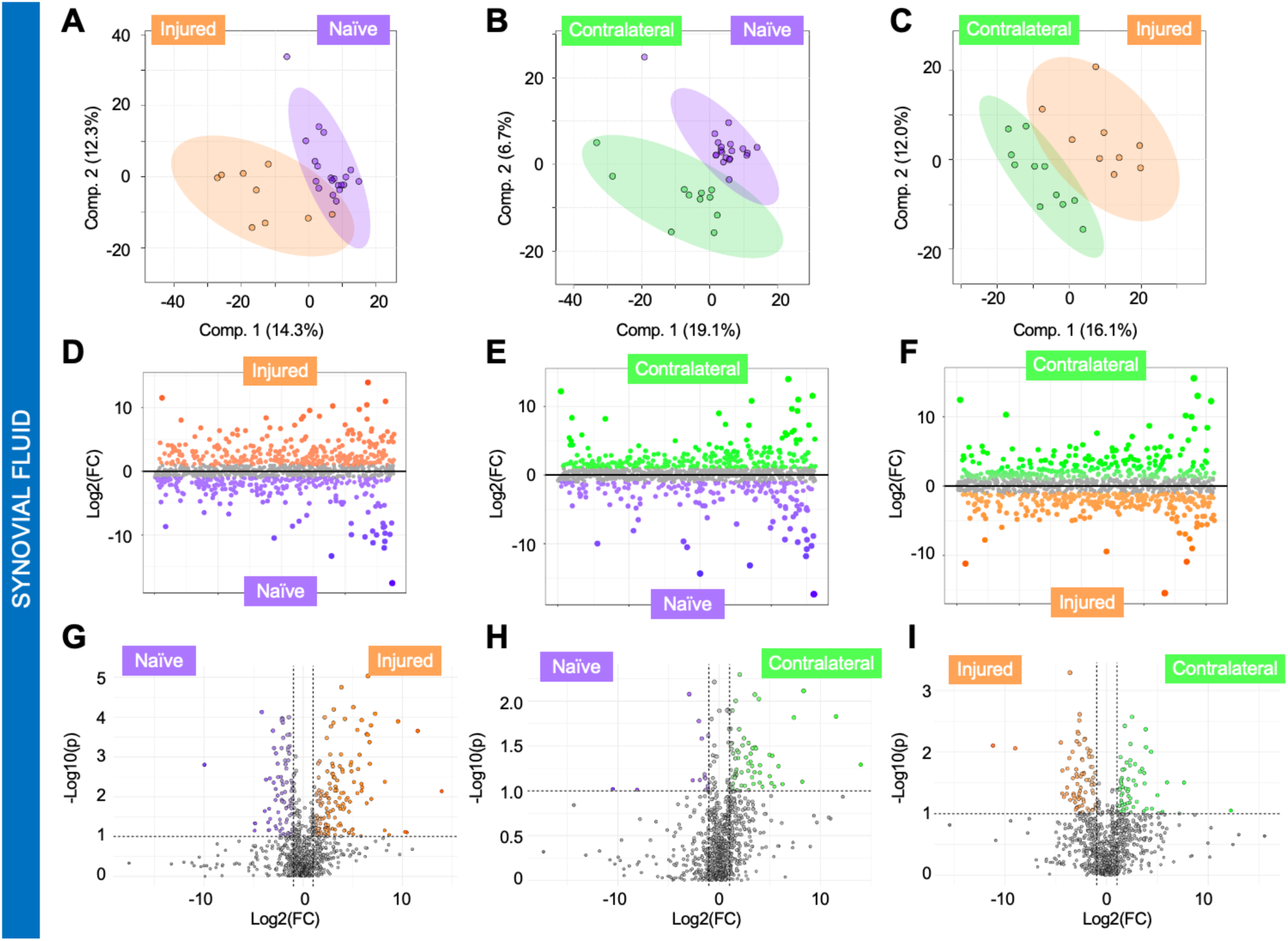
Synovial fluid metabolome differs across injured, contralateral, and naïve limbs. (A-C) Partial Least Squares-Discriminant Analysis (PLS-DA) finds some overlap between (A) injured and naïve, (B) naïve and contralateral, and (C) injured and contralateral synovial fluid. (D-F) Fold change analysis distinguished populations of metabolite features driving separation of metabolomic profiles. (D) Specifically, 318 and 253 metabolite features were highest in injured and naïve synovial fluid, respectively. (E) 269 and 178 metabolite features were highest in contralateral and naïve synovial fluid, respectively. (F) 303 and 291 features were highest in contralateral and injured synovial fluid, respectively. (G-I) To further examine metabolic differences associated with injury status across whole joints, volcano plot analysis was performed and identified numerous metabolite features that had a fold change > 2, a p-value < 0.05, and were differentially regulated between injured and naïve (G, n = 59), contralateral and naive (H, n = 81), and contralateral and injured synovial fluid (I, n = 140). The colors in A-I correspond to: purple = naïve synovial fluid, orange = injured synovial fluid, green = contralateral synovial fluid.

**Figure S5.**
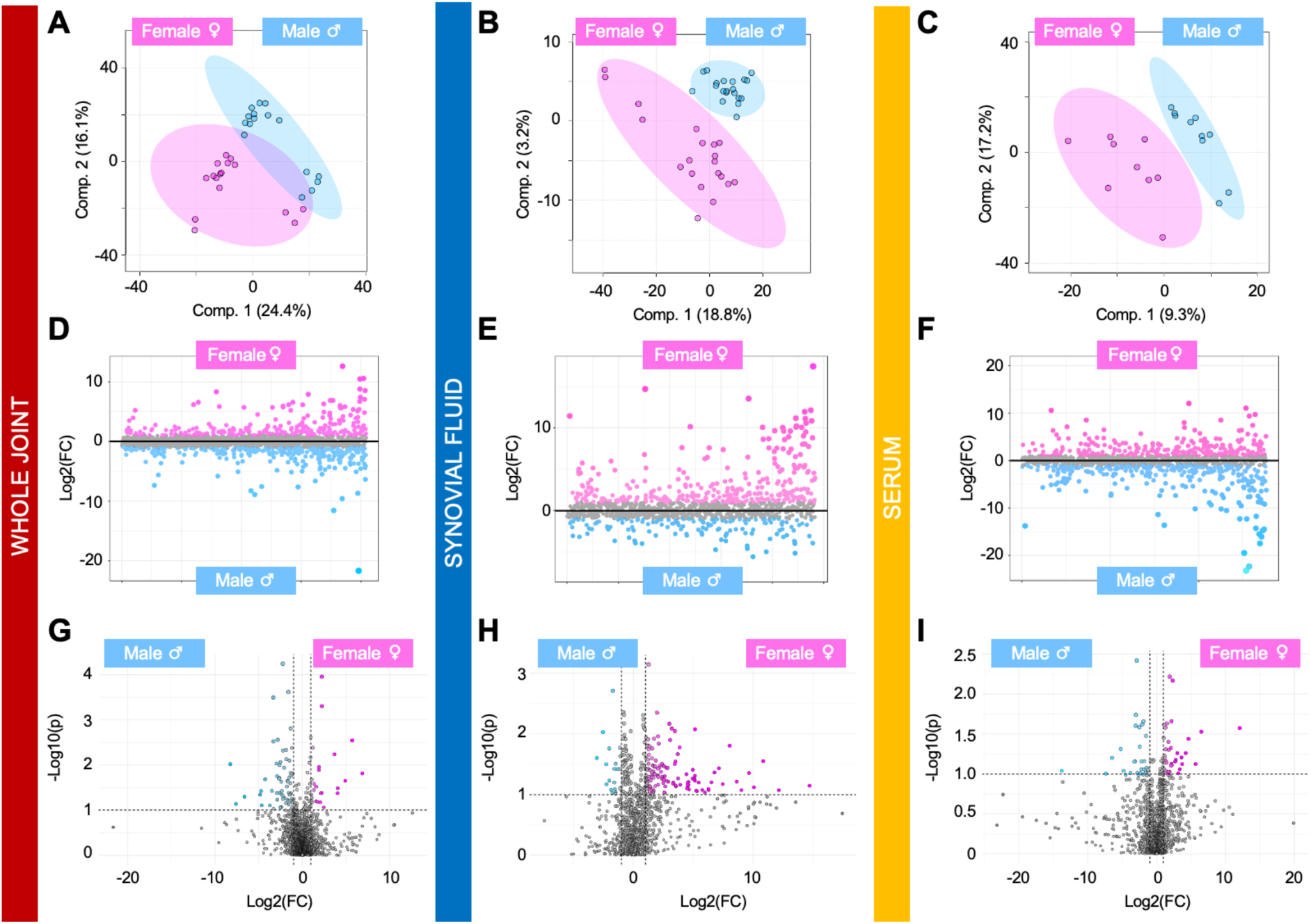
Sexual dimorphic patterns are detected across whole joints, synovial fluid, and serum from injured and naïve mice. (A-C) Partial Least Squares-Discriminant Analysis (PLS-DA) finds (A) minimal overlap of male and female whole joints, while (B) synovial fluid and (C) serum from males and females perfectly clusters apart from each other. (D-F) Fold change analysis distinguished populations of metabolite features driving separation of male and female metabolomic profiles. (D) Specifically, 253 and 314 metabolite features were highest in female and male whole joints, respectively. (E) 334 and 137 metabolite features were highest in female and male synovial fluid, respectively. (F) 267 and 315 features were highest in female and male serum, respectively. (G-I) To further examine metabolic differences associated with sex across tissue types, volcano plot analysis was performed and identified numerous metabolite features that had a fold change > 2, a p-value < 0.05, and were differentially regulated between male and female whole joints (G, n = 90), synovial fluid (H, n = 159), and serum (I, n = 63). The colors in A-I correspond to: pink = females, blue = males, red = whole joint, blue = synovial fluid, yellow = serum.

**Figure S6.**
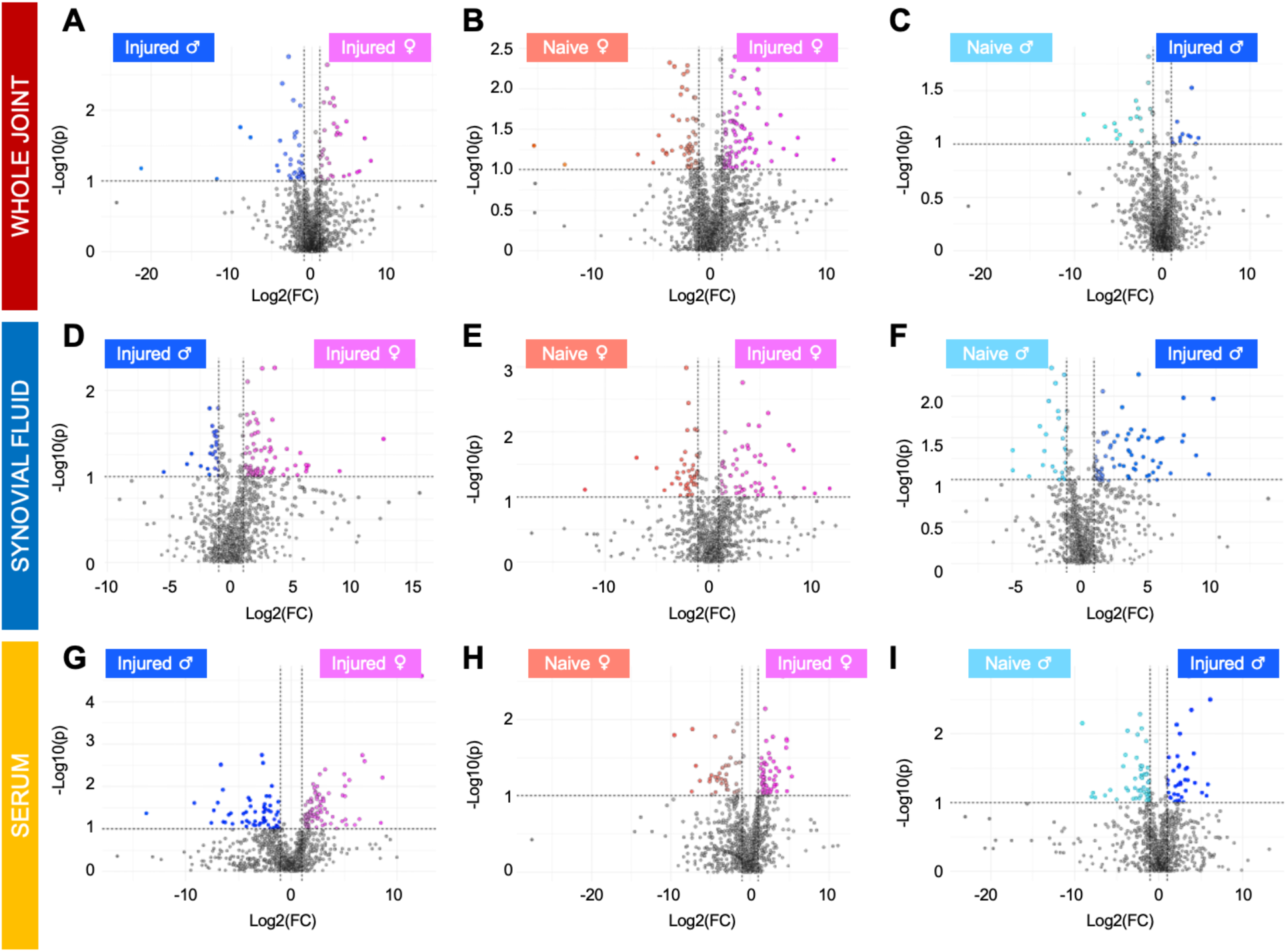
Volcano plot analysis reveals both sex- and injury-associated metabolites across sample types. (A-C) To pinpoint metabolic differences associated with injury and sex at the whole joint level, volcano plot analysis was performed and identified numerous metabolite features that had a fold change > 2, a p-value < 0.05, and were differentially regulated between injured males and females (A, n = 110), injured and naïve females (B, n = 158), and injured and naïve males (C, n = 32). (D-F) Among SF samples, volcano plot analysis identified metabolites that were differentially regulated between injured males and females (D, n = 95), injured and naïve females (E, n = 101), and injured and naïve males (F, n = 99). (G-I) At the serum level, a similar pattern was found where volcano plot analysis identified metabolites among injured males and females (G, n = 82), injured and naïve females (H, n = 92), and injured and naïve males (I, n = 87). The colors in A-I correspond to: pink = injured females, peach = naïve females, royal blue = injured males, light blue = naïve males, red = whole joint, blue = synovial fluid, yellow = serum.

**Figure S7.**
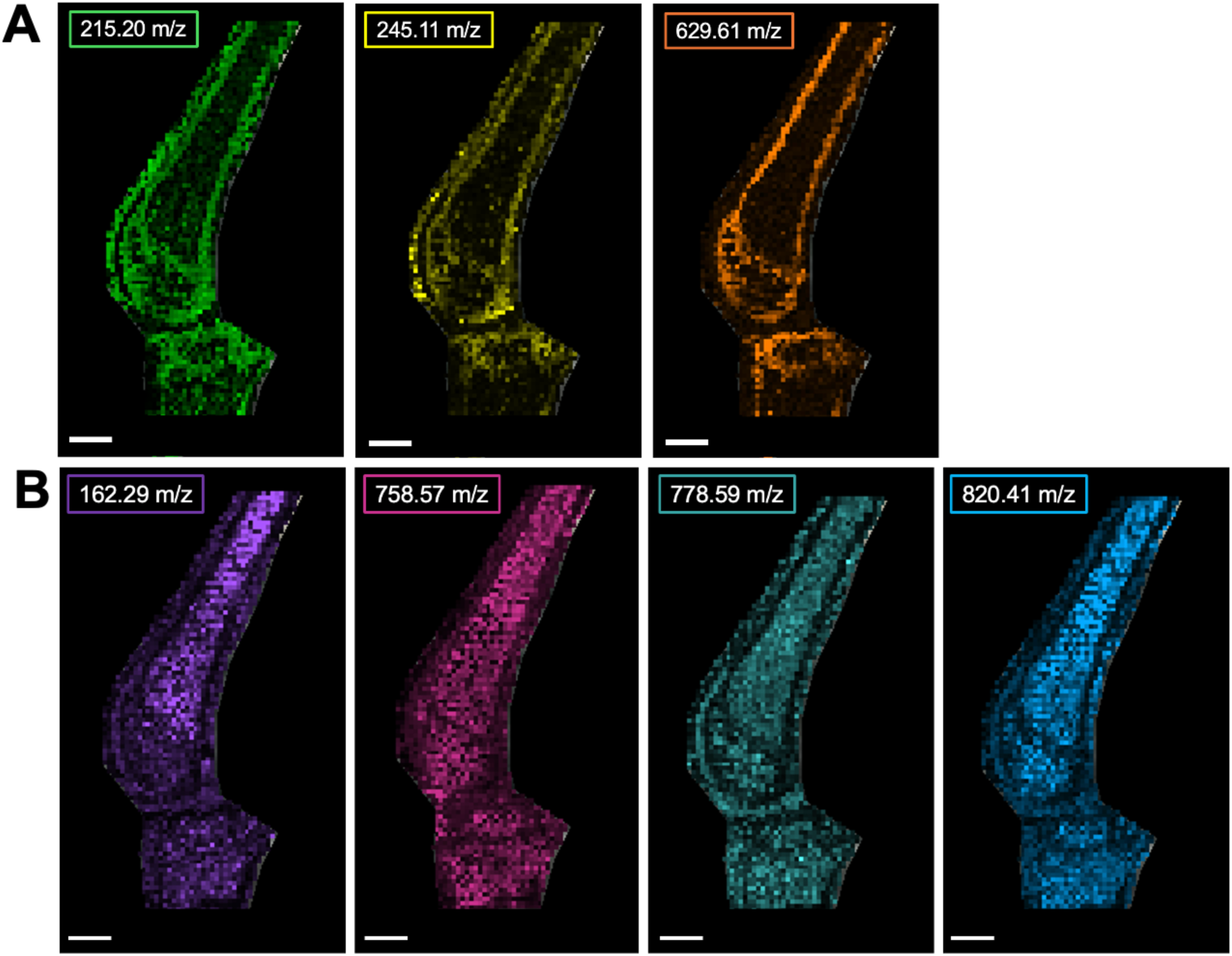
MALDI-MSI analysis combined with LC-MS/MS detects and identifies differences in osteochondral metabolites. (A) Molecular species that were putatively identified with notable spatial patterns amongst bone and the growth plate include alpha-carboxy-delta-decalactone (215.20, green), hydroxyprolyl-isoleucine (245.11 m/z, yellow), and C_36_H_38_O_7_ (629.61 m/z, orange). (B) conversely, molecular species that were putatively identified with notable patterns among bone marrow regions include L-carnitine (162.29 m/z, purple) and various lipid species (34 carbons, 2 double bonds – 758.57 m/z, pink; 34 double bonds, 3 double bonds – 778.59 m/z, teal; 39 carbons, 6 double bonds – 820.41 m/z, blue). Spatial resolution = 100 um. Scale bar = 1 mm. Interval width = 0.35 Da.

**Figure S8.**
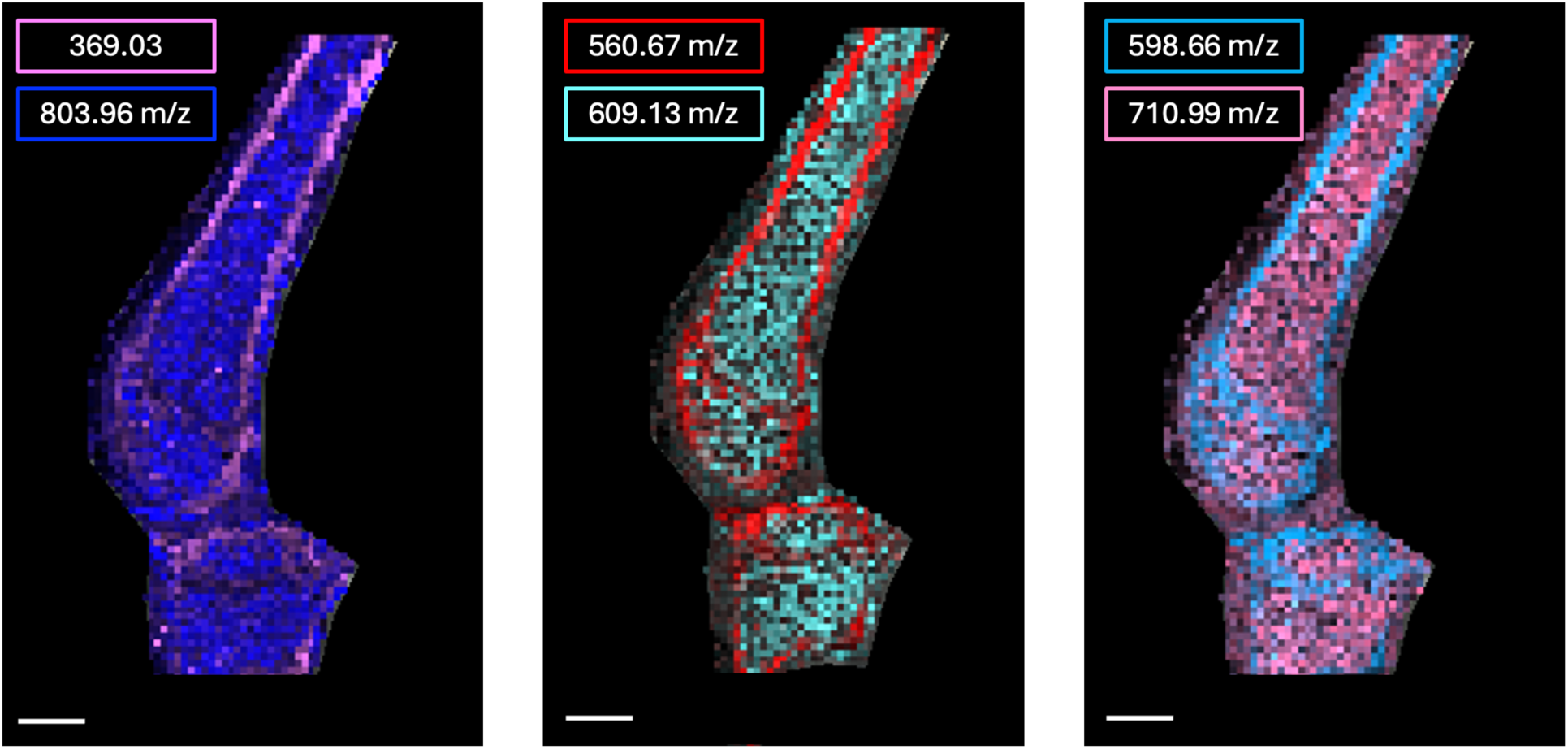
Unidentified molecular species localize to different structures within the joint. Species more abundant in bone and growth plate included 369.03 m/z (pink), 560.67 m/z (red), and 598.66 m/z (light blue). Conversely, those more abundant in bone marrow included 803.96 m/z (dark blue), 609.13 m/z (cyan), and 710.99 m/z (light pink). Spatial resolution = 100 um. Scale bar = 1 mm. Interval width = 0.35 Da.

**Figure S9.**
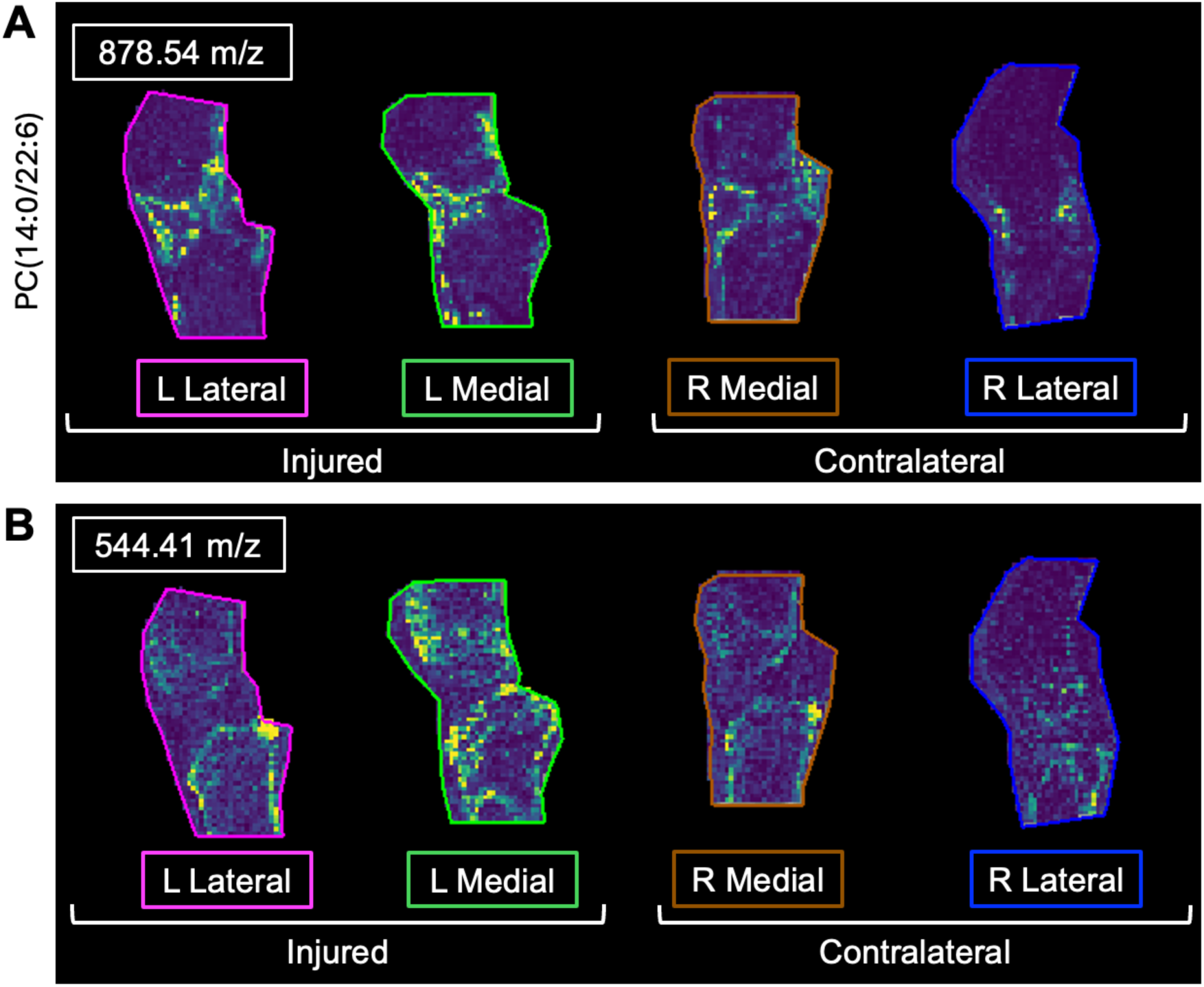
Injury-associated spatial distribution patterns between injury and contralateral limbs. (A) 544.41 m/z and (B) putatively identified PC(14:0/22:6) (878.54 m/z) display notable injury associated patterns between medial and lateral sagittal sections from an injured and contralateral joint. Left to right – left lateral, left medial, right medial, right lateral. Spatial resolution = 100 um. Scale bar = 1 mm. Interval width = 0.35 Da.

**Figure S10.**
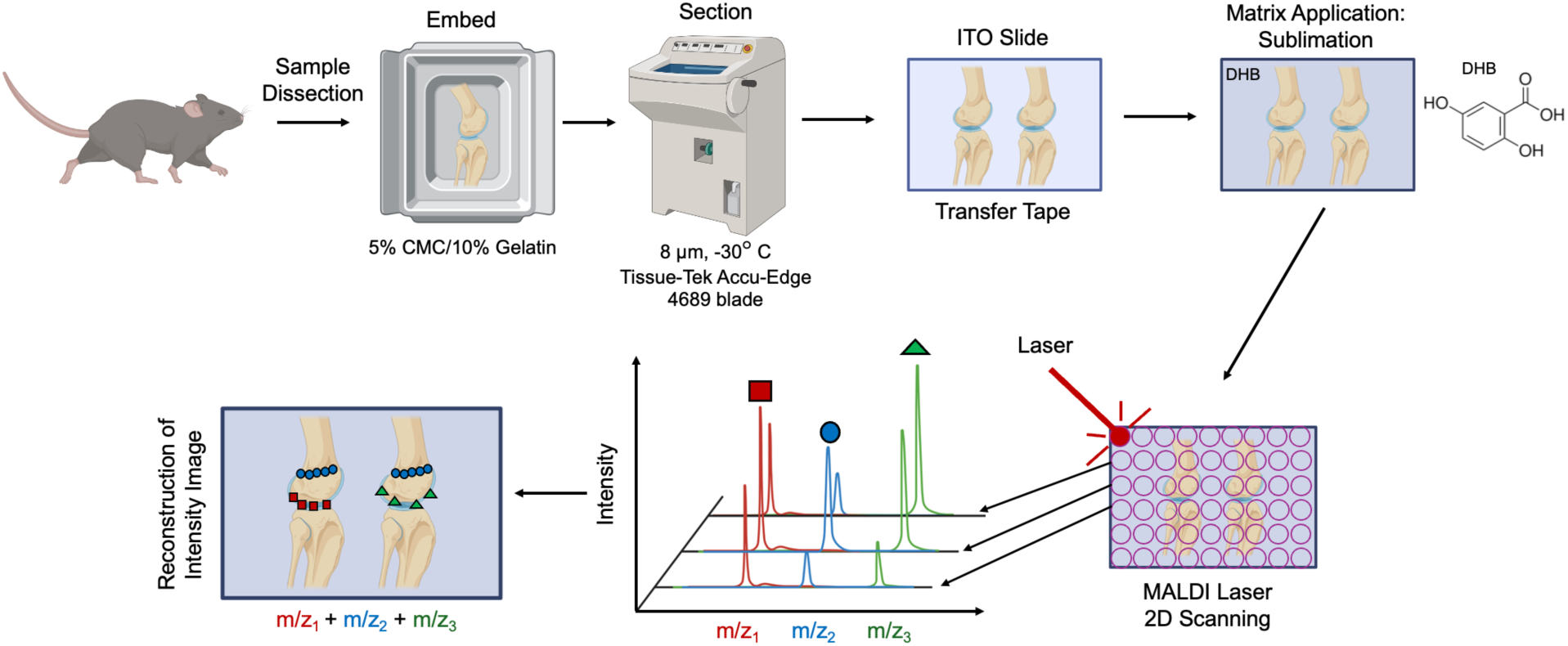
MALDI-MSI experimental workflow to spatially image osteochondral metabolites. Whole joints were obtained from C57Bl6/J male and female injured and naïve mice, embedded in 5% carboxylmethylcellulose (CMC)/10% gelatin, sectioned (8 um), transferred to indium tin oxide (ITO) slides, and sublimed with 2,5-dihydroxybenzoic acid matrix. Data were then acquired using 2D laser scanning followed by reconstruction of intensity images representing m/z values and intensity.

**Figure S11.**
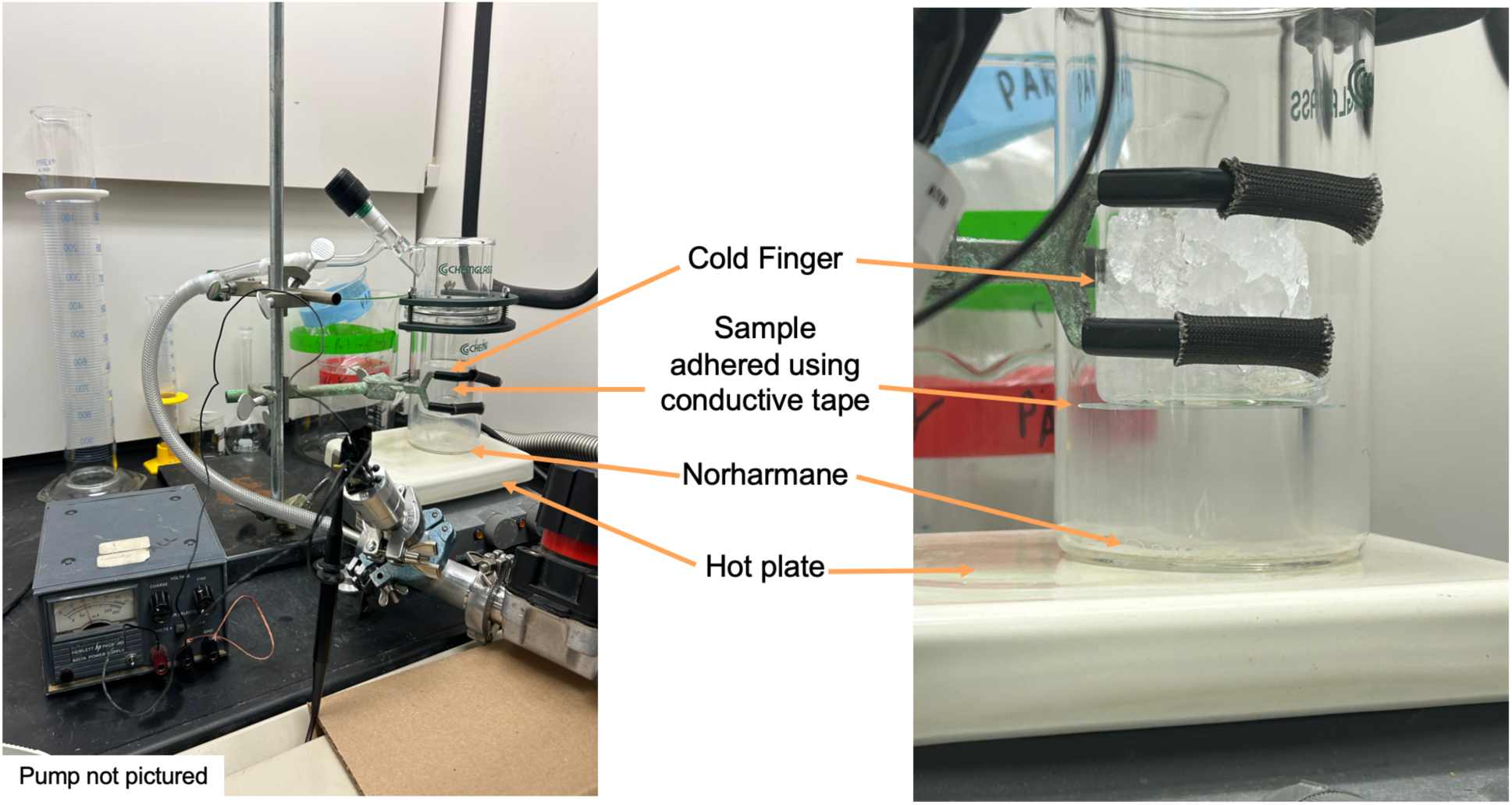
Sublimation apparatus used to uniformly coat whole joints with matrix. Components of the sublimation apparatus include the cold finger, hot plate, glass condenser and sleeve, O-ring seal, and vacuum pump (not pictured).

## References

[1] T.D. Brown, R.C. Johnston, C.L. Saltzman, J.L. Marsh, J.A. Buckwalter, Posttraumatic osteoarthritis: a first estimate of incidence, prevalence, and burden of disease, J Orthop Trauma 20(10) (2006) 739–44.

[2] A.C. Thomas, T. Hubbard-Turner, E.A. Wikstrom, R.M. Palmieri-Smith, Epidemiology of Posttraumatic Osteoarthritis, J Athl Train 52(6) (2017) 491–496.

[3] C.A. Gottlob, C.L. Baker, Jr., J.M. Pellissier, L. Colvin, Cost effectiveness of anterior cruciate ligament reconstruction in young adults, Clin Orthop Relat Res (367) (1999) 272–82.

[4] L.Y. Griffin, M.J. Albohm, E.A. Arendt, R. Bahr, B.D. Beynnon, M. Demaio, R.W. Dick, L. Engebretsen, W.E. Garrett, Jr., J.A. Hannafin, T.E. Hewett, L.J. Huston, M.L. Ireland, R.J. Johnson, S. Lephart, B.R. Mandelbaum, B.J. Mann, P.H. Marks, S.W. Marshall, G. Myklebust, F.R. Noyes, C. Powers, C. Shields, Jr., S.J. Shultz, H. Silvers, J. Slauterbeck, D.C. Taylor, C.C. Teitz, E.M. Wojtys, B. Yu, Understanding and preventing noncontact anterior cruciate ligament injuries: a review of the Hunt Valley II meeting, January 2005, Am J Sports Med 34(9) (2006) 1512–32.

[5] L.S. Lohmander, P.M. Englund, L.L. Dahl, E.M. Roos, The long-term consequence of anterior cruciate ligament and meniscus injuries: osteoarthritis, Am J Sports Med 35(10) (2007) 1756–69.

[6] M.E. Cinque, G.J. Dornan, J. Chahla, G. Moatshe, R.F. LaPrade, High Rates of Osteoarthritis Develop After Anterior Cruciate Ligament Surgery: An Analysis of 4108 Patients, Am J Sports Med 46(8) (2018) 2011–2019.

[7] D.N. Giugliano, J.L. Solomon, ACL tears in female athletes, Phys Med Rehabil Clin N Am 18(3) (2007) 417–38, viii.

[8] A.M. Bruder, A.G. Culvenor, M.G. King, M. Haberfield, E.A. Roughead, J. Mastwyk, J.L. Kemp, M. Ferraz Pazzinatto, T.J. West, S.L. Coburn, S.M. Cowan, A.M. Ezzat, L. To, K. Chilman, J.L. Couch, J.L. Whittaker, K.M. Crossley, Let’s talk about sex (and gender) after ACL injury: a systematic review and meta-analysis of self-reported activity and knee-related outcomes, Br J Sports Med 57(10) (2023) 602–610.

[9] C.L. Blaker, D.M. Ashton, N. Doran, C.B. Little, E.C. Clarke, Sex- and injury-based differences in knee biomechanics in mouse models of post-traumatic osteoarthritis, J Biomech 114 (2021) 110152.

[10] V.K. Srikanth, J.L. Fryer, G. Zhai, T.M. Winzenberg, D. Hosmer, G. Jones, A meta-analysis of sex differences prevalence, incidence and severity of osteoarthritis, Osteoarthritis Cartilage 13(9) (2005) 769–81.

[11] S.C. Faber, F. Eckstein, S. Lukasz, R. Muhlbauer, J. Hohe, K.H. Englmeier, M. Reiser, Gender differences in knee joint cartilage thickness, volume and articular surface areas: assessment with quantitative three-dimensional MR imaging, Skeletal Radiol 30(3) (2001) 144–50.

[12] K. Hitt, J.R. Shurman, 2nd, K. Greene, J. McCarthy, J. Moskal, T. Hoeman, M.A. Mont, Anthropometric measurements of the human knee: correlation to the sizing of current knee arthroplasty systems, J Bone Joint Surg Am 85-A Suppl 4 (2003) 115-22.

[13] B.D. Hislop, C. Devine, R.K. June, C.M. Heveran, Subchondral bone structure and synovial fluid metabolism are altered in injured and contralateral limbs 7 days after non-invasive joint injury in skeletally-mature C57BL/6 mice, Osteoarthritis Cartilage (2022).

[14] B. Mickiewicz, B.J. Heard, J.K. Chau, M. Chung, D.A. Hart, N.G. Shrive, C.B. Frank, H.J. Vogel, Metabolic profiling of synovial fluid in a unilateral ovine model of anterior cruciate ligament reconstruction of the knee suggests biomarkers for early osteoarthritis, J Orthop Res 33(1) (2015) 71–7.

[15] C.W. Wallace, B. Hislop, A.K. Hahn, A.E. Erdogan, P.P. Brahmachary, R.K. June, Correlations between metabolites in the synovial fluid and serum: A mouse injury study, J Orthop Res (2022).

[16] H.D. Welhaven, A.H. Welfley, P. Pershad, J. Satalich, R. O’Connell, B. Bothner, A.R. Vap, R.K. June, Metabolic phenotypes reflect patient sex and injury status: A cross-sectional analysis of human synovial fluid, Osteoarthritis Cartilage (2023).

[17] A.K. Hahn, C.W. Wallace, H.D. Welhaven, E. Brooks, M. McAlpine, B.A. Christiansen, S.T. Walk, R.K. June, The microbiome mediates epiphyseal bone loss and metabolomic changes after acute joint trauma in mice, Osteoarthritis Cartilage 29(6) (2021) 882–893.

[18] E.H. Seeley, R.M. Caprioli, Molecular imaging of proteins in tissues by mass spectrometry, Proc Natl Acad Sci U S A 105(47) (2008) 18126–31.

[19] E.R. Amstalden van Hove, D.F. Smith, R.M. Heeren, A concise review of mass spectrometry imaging, J Chromatogr A 1217(25) (2010) 3946–54.

[20] B. Cillero-Pastor, G.B. Eijkel, A. Kiss, F.J. Blanco, R.M. Heeren, Matrix-assisted laser desorption ionization-imaging mass spectrometry: a new methodology to study human osteoarthritic cartilage, Arthritis Rheum 65(3) (2013) 710–20.

[21] B. Cillero-Pastor, G. Eijkel, A. Kiss, F.J. Blanco, R.M. Heeren, Time-of-flight secondary ion mass spectrometry-based molecular distribution distinguishing healthy and osteoarthritic human cartilage, Anal Chem 84(21) (2012) 8909–16.

[22] M.J. Peffers, B. Cillero-Pastor, G.B. Eijkel, P.D. Clegg, R.M. Heeren, Matrix assisted laser desorption ionization mass spectrometry imaging identifies markers of ageing and osteoarthritic cartilage, Arthritis Res Ther 16(3) (2014) R110.

[23] B. Cillero-Pastor, G.B. Eijkel, F.J. Blanco, R.M. Heeren, Protein classification and distribution in osteoarthritic human synovial tissue by matrix-assisted laser desorption ionization mass spectrometry imaging, Anal Bioanal Chem 407(8) (2015) 2213–22.

[24] C.J. Good, E.K. Neumann, C.E. Butrico, J.E. Cassat, R.M. Caprioli, J.M. Spraggins, High Spatial Resolution MALDI Imaging Mass Spectrometry of Fresh-Frozen Bone, Anal Chem 94(7) (2022) 3165–3172.

[25] M. Yamauchi, M. Terajima, M. Shiiba, Lysine Hydroxylation and Cross-Linking of Collagen, Methods Mol Biol 1934 (2019) 309–324.

[26] S. Pornprasertsuk, W.R. Duarte, Y. Mochida, M. Yamauchi, Overexpression of lysyl hydroxylase-2b leads to defective collagen fibrillogenesis and matrix mineralization, J Bone Miner Res 20(1) (2005) 81–7.

[27] M. Saito, K. Marumo, Collagen cross-links as a determinant of bone quality: a possible explanation for bone fragility in aging, osteoporosis, and diabetes mellitus, Osteoporos Int 21(2) (2010) 195–214.

[28] A.K. Carlson, R.A. Rawle, C.W. Wallace, E.G. Brooks, E. Adams, M.C. Greenwood, M. Olmer, M.K. Lotz, B. Bothner, R.K. June, Characterization of synovial fluid metabolomic phenotypes of cartilage morphological changes associated with osteoarthritis, Osteoarthritis Cartilage 27(8) (2019) 1174–1184.

[29] K. Tootsi, K. Vilba, A. Martson, J. Kals, K. Paapstel, M. Zilmer, Metabolomic Signature of Amino Acids, Biogenic Amines and Lipids in Blood Serum of Patients with Severe Osteoarthritis, Metabolites 10(8) (2020).

[30] A. Ohnishi, T. Osaki, Y. Matahira, T. Tsuka, T. Imagawa, Y. Okamoto, S. Minami, Correlation of plasma amino acid concentrations and chondroprotective effects of glucosamine and fish collagen peptide on the development of osteoarthritis, J Vet Med Sci 75(4) (2013) 497–502.

[31] Y. Hu, Q. Wu, Y. Qiao, P. Zhang, W. Dai, H. Tao, S. Chen, Disturbances in Metabolic Pathways and the Identification of a Potential Biomarker Panel for Early Cartilage Degeneration in a Rabbit Anterior Cruciate Ligament Transection Model, Cartilage 13(2_suppl) (2021) 1376S-1387S.

[32] H.D. Welhaven, A.H. Welfley, P. Pershad, J. Satalich, R. O’Connell, B. Bothner, A.R. Vap, R.K. June, Metabolic phenotypes reflect patient sex and injury status: A cross-sectional analysis of human synovial fluid, Osteoarthritis Cartilage 32(9) (2024) 1074–1083.

[33] K.H. Ding, M. Cain, M. Davis, C. Bergson, M. McGee-Lawrence, C. Perkins, T. Hardigan, X. Shi, Q. Zhong, J. Xu, W.B. Bollag, W. Hill, M. Elsalanty, M. Hunter, M.C. Isales, P. Lopez, M. Hamrick, C.M. Isales, Amino acids as signaling molecules modulating bone turnover, Bone 115 (2018) 15–24.

[34] A.D. Conigrave, E.M. Brown, R. Rizzoli, Dietary protein and bone health: roles of amino acid-sensing receptors in the control of calcium metabolism and bone homeostasis, Annu Rev Nutr 28 (2008) 131–55.

[35] G. Yang, H. Zhang, T. Chen, W. Zhu, S. Ding, K. Xu, Z. Xu, Y. Guo, J. Zhang, Metabolic analysis of osteoarthritis subchondral bone based on UPLC/Q-TOF-MS, Anal Bioanal Chem 408(16) (2016) 4275–86.

[36] T. Igari, M. Tsuchizawa, T. Shimamura, Alteration of tryptophan metabolism in the synovial fluid of patients with rheumatoid arthritis and osteoarthritis, Tohoku J Exp Med 153(2) (1987) 79–86.

[37] K.Y. Kang, S.H. Lee, S.M. Jung, S.H. Park, B.H. Jung, J.H. Ju, Downregulation of Tryptophan-related Metabolomic Profile in Rheumatoid Arthritis Synovial Fluid, J Rheumatol 42(11) (2015) 2003–11.

[38] E.M. Leimer, L.M. Tanenbaum, D.L. Nettles, R.D. Bell, M.E. Easley, L.A. Setton, S.B. Adams, Amino Acid Profile of Synovial Fluid Following Intra-articular Ankle Fracture, Foot Ankle Int 39(10) (2018) 1169–1177.

[39] H. Akasaka, H. Yoshida, H. Takizawa, N. Hanawa, T. Tobisawa, M. Tanaka, N. Moniwa, N. Togashi, T. Yamashita, S. Kuroda, N. Ura, T. Miura, B.-C. Investigators, The impact of elevation of serum uric acid level on the natural history of glomerular filtration rate (GFR) and its sex difference, Nephrol Dial Transplant 29(10) (2014) 1932–9.

[40] S.L. Mumford, S.S. Dasharathy, A.Z. Pollack, N.J. Perkins, D.R. Mattison, S.R. Cole, J. Wactawski-Wende, E.F. Schisterman, Serum uric acid in relation to endogenous reproductive hormones during the menstrual cycle: findings from the BioCycle study, Hum Reprod 28(7) (2013) 1853–62.

[41] G.R. Steinberg, S.L. Macaulay, M.A. Febbraio, B.E. Kemp, AMP-activated protein kinase--the fat controller of the energy railroad, Can J Physiol Pharmacol 84(7) (2006) 655–65.

[42] J. Wang, J. Li, D. Song, J. Ni, M. Ding, J. Huang, M. Yan, AMPK: implications in osteoarthritis and therapeutic targets, Am J Transl Res 12(12) (2020) 7670–7681.

[43] D. Yi, H. Yu, K. Lu, C. Ruan, C. Ding, L. Tong, X. Zhao, D. Chen, AMPK Signaling in Energy Control, Cartilage Biology, and Osteoarthritis, Front Cell Dev Biol 9 (2021) 696602.

[44] T. Purdom, L. Kravitz, K. Dokladny, C. Mermier, Understanding the factors that effect maximal fat oxidation, J Int Soc Sports Nutr 15 (2018) 3.

[45] S. Yang, J. Wang, Estrogen Activates AMP-Activated Protein Kinase in Human Endothelial Cells via ERbeta/Ca(2+)/Calmodulin-Dependent Protein Kinase Kinase beta Pathway, Cell Biochem Biophys 72(3) (2015) 701–7.

[46] N. Cui, M. Hu, R.A. Khalil, Biochemical and Biological Attributes of Matrix Metalloproteinases, Prog Mol Biol Transl Sci 147 (2017) 1–73.

[47] R. Visse, H. Nagase, Matrix metalloproteinases and tissue inhibitors of metalloproteinases: structure, function, and biochemistry, Circ Res 92(8) (2003) 827–39.

[48] R.P. Verma, C. Hansch, Matrix metalloproteinases (MMPs): chemical-biological functions and (Q)SARs, Bioorg Med Chem 15(6) (2007) 2223–68.

[49] Y.J. Lee, E.B. Lee, Y.E. Kwon, J.J. Lee, W.S. Cho, H.A. Kim, Y.W. Song, Effect of estrogen on the expression of matrix metalloproteinase (MMP)-1, MMP-3, and MMP-13 and tissue inhibitor of metalloproternase-1 in osteoarthritis chondrocytes, Rheumatol Int 23(6) (2003) 282–8.

[50] A.K. Carlson, R.A. Rawle, E. Adams, M.C. Greenwood, B. Bothner, R.K. June, Application of global metabolomic profiling of synovial fluid for osteoarthritis biomarkers, Biochem Biophys Res Commun 499(2) (2018) 182–188.

[51] M.R. Eveque-Mourroux, P.J. Emans, A. Boonen, B.S.R. Claes, F.G. Bouwman, R.M.A. Heeren, B. Cillero-Pastor, Heterogeneity of Lipid and Protein Cartilage Profiles Associated with Human Osteoarthritis with or without Type 2 Diabetes Mellitus, J Proteome Res 20(5) (2021) 2973–2982.

[52] P. Pousinis, P.R.W. Gowler, J.J. Burston, C.A. Ortori, V. Chapman, D.A. Barrett, Lipidomic identification of plasma lipids associated with pain behaviour and pathology in a mouse model of osteoarthritis, Metabolomics 16(3) (2020) 32.

[53] A.K. Carlson, R.A. Rawle, C.W. Wallace, E. Adams, M.C. Greenwood, B. Bothner, R.K. June, Global metabolomic profiling of human synovial fluid for rheumatoid arthritis biomarkers, Clin Exp Rheumatol 37(3) (2019) 393–399.

[54] H.D. Welhaven, E. Viles, J. Starke, C. Wallace, B. Bothner, R.K. June, A.K. Hahn, Metabolomic profiles of cartilage and bone reflect tissue type, radiography-confirmed osteoarthritis, and spatial location within the joint, Biochem Biophys Res Commun 703 (2024) 149683.

[55] H.D. Welhaven, A.H. Welfley, P. Brahmachary, A.R. Bergstrom, E. Houske, M. Glimm, B. Bothner, A.K. Hahn, R.K. June, Metabolomic Profiles and Pathways in Osteoarthritic Human Cartilage: A Comparative Analysis with Healthy Cartilage, Metabolites 14(4) (2024).

[56] B.A. Christiansen, M.J. Anderson, C.A. Lee, J.C. Williams, J.H. Yik, D.R. Haudenschild, Musculoskeletal changes following non-invasive knee injury using a novel mouse model of post-traumatic osteoarthritis, Osteoarthritis Cartilage 20(7) (2012) 773–82.

[57] B.A. Christiansen, F. Guilak, K.A. Lockwood, S.A. Olson, A.A. Pitsillides, L.J. Sandell, M.J. Silva, M.C. van der Meulen, D.R. Haudenschild, Non-invasive mouse models of post-traumatic osteoarthritis, Osteoarthritis Cartilage 23(10) (2015) 1627–38.

[58] C.B. Little, D.J. Hunter, Post-traumatic osteoarthritis: from mouse models to clinical trials, Nat Rev Rheumatol 9(8) (2013) 485–97.

[59] C.W. Wallace, B. Hislop, A.K. Hahn, A.E. Erdogan, P.P. Brahmachary, R.K. June, Correlations between metabolites in the synovial fluid and serum: A mouse injury study, J Orthop Res 40(12) (2022) 2792–2802.

[60] D.R. Seifer, B.D. Furman, F. Guilak, S.A. Olson, S.C. Brooks, 3rd, V.B. Kraus, Novel synovial fluid recovery method allows for quantification of a marker of arthritis in mice, Osteoarthritis Cartilage 16(12) (2008) 1532–8.

[61] Z. Pang, G. Zhou, J. Ewald, L. Chang, O. Hacariz, N. Basu, J. Xia, Using MetaboAnalyst 5.0 for LC-HRMS spectra processing, multi-omics integration and covariate adjustment of global metabolomics data, Nat Protoc 17(8) (2022) 1735–1761.

[62] K.A. Nelson, G.J. Daniels, J.W. Fournie, M.J. Hemmer, Optimization of whole-body zebrafish sectioning methods for mass spectrometry imaging, J Biomol Tech 24(3) (2013) 119–27.

[63] T. Kawamoto, K. Kawamoto, Preparation of thin frozen sections from nonfixed and undecalcified hard tissues using Kawamot’s film method (2012), Methods Mol Biol 1130 (2014) 149–164.

